# Motor sequence learning involves better prediction of the next action and optimization of movement trajectories

**DOI:** 10.1101/2024.12.23.630092

**Authors:** Mehrdad Kashefi, Jörn Diedrichsen, J. Andrew Pruszynski

## Abstract

Learning new sequential movements is a fundamental skill for many animals. Although the behavioral manifestations of sequence learning are clear, the underlying mechanisms remain poorly understood. Motor sequence learning may arise from three distinct processes: (1) improved execution of individual movements independent of their sequential context; (2) enhanced anticipation of “what” movement should be executed next, enabling faster initiation; and (3) the development of motoric sequence-specific representations that encode “how” movements should be optimally performed within a sequence. However, many existing paradigms conflate the "what" and "how" components of learning, as participants often acquire both the sequence content (what to do) and its execution (how to do it). This overlap obscures the distinct contributions of each mechanism to motor sequence learning. In this study, we disentangled these mechanisms using a continuous reaching task. Performance in trained sequences was compared to random sequences to rule out improvements attributable solely to isolated movement execution. By also varying how many upcoming targets were visible we assessed the role of anticipation in learning. When participants could only see one future target, improvements were mostly due to them learning which target would come next. When they could see four future targets, participants immediately demonstrated fast movement times and increased movement smoothness, surpassing late-stage performance in the one target condition. Crucially, even with full visibility of future targets, participants showed further sequence-specific learning caused by a continuous optimization of movement trajectories. Follow-up experiments revealed that the learned sequence representations were effector-specific and encoded contextual information of four movements or longer. Our paradigm enables a clear dissociation between the "what" and "how" components of motor sequence learning and provides compelling evidence for the development of effector-specific sequence representations that guide optimal movement execution.

## Introduction

Many animals can expand their motor repertoire by learning new movement sequences – with songbirds (Fee and Scharff, 2010) and pianists (Engel et al., 1997) being exceptional examples of this ability. The behavioral markers of sequence learning are clear: with practice, motor sequences are performed more accurately, quickly, smoothly, (Berlot et al., 2020; Karni et al., 1998; Moisello et al., 2009) and with less cognitive effort (Keele et al., 2003; Pashler, 1994; Reber and Squire, 1994). The learning mechanisms that underlie these improvements, however, remain elusive (Diedrichsen and Kornysheva, 2015; Krakauer et al., 2019; Warren et al., 2011; Wong and Krakauer, 2019).

Multiple mechanisms can contribute to improvements in motor sequence tasks. First, practice can refine the individual movements, independent of their sequential context. Second, sequence learning can reflect getting better at knowing *what* to do next (Perruchet and Amorim, 1992) by learning to anticipate the location of the next target in a sequential reaching task or which key to press in finger sequencing task. Finally, sequence learning may reflect the acquisition of a motoric sequence representation, which specifies *how* to execute a specific sequence skillfully (Karni et al., 1998, 1995; Picard et al., 2013)

One paradigm that has been extensively used to study motor sequence learning is the Serial Reaction Time Task (SRTT; Nissen and Bullemer, 1987). The SRTT involves responding to visual stimuli that are presented one by one on a screen. Unbeknownst to the participants, these stimuli can follow a specific sequence and, after many repetitions of the specific sequence, participant reaction time decreases. Based on the observation that learning in the SRTT can occur outside of conscious awareness, improvement has been attributed to the formation of a procedural memory (Nissen et al., 1989; Nissen and Bullemer, 1987; Reber and Squire, 1998; Willingham et al., 1989). However it has been argued that most sequence-specific learning in this task arises because participants get better at anticipating *what* to do next (Willingham, 1999; Willingham et al., 2000). In fact, Wong et al. (2015) show that when the SRTT is designed such that participants can fully predict the next stimulus, practicing the sequence does not lead to any sequence-specific improvements.

This observation raises the question whether improvements in other motor sequence learning tasks are also attributable to better anticipation. In many motor sequence paradigms participants have at least partial knowledge of the upcoming movement elements—either because they are explicitly taught the sequence or because the sequence items are presented visually in the form of digits on a screen (Berlot et al., 2020; Karni et al., 1995; Verwey, 2003; Verwey and Wright, 2004; Yokoi et al., 2017). Do participants in this case only improve how quickly they can decide *what* key to press, or do they learn and optimize a motoric sequence representation?

Here we addressed this question with a sequence paradigm in which participants reached continuously to visually presented targets. Participants either saw only the next target on the screen (Horizon 1, effectively a version of the SRTT), or they saw the next four targets on the screen (Horizon 4). The sequences were either repeatedly practiced or different from trial to trial. We showed that most of the learning in SRTT conditions was merely learning to anticipate the location of the next target. We did so by comparing sequence learning with and without foreknowledge of future reach targets. However, even with foreknowledge of future reach targets, participants improved their performance further. In two follow-up experiments, we further showed that this sequence-specific representation was effector-specific, and that it generalizes to other sequences that had elements of 4 or more reaches in common with the originally learned sequence.

## Results

We trained participants to perform a sequential reaching task in an exoskeleton robot. Participants moved a cursor that veridically represented the position of their right hand. On each trial, participants had to move the cursor to a set of 14 circular targets as quickly and accurately as possible (**Fig. 1A**, **see Methods**). Target locations were selected with a rule that forced consecutive targets to be 5 cm apart. We measured sequence-specific learning by contrasting participants’ performance between a set of sequences that were executed once (**Random**) and a single fixed sequence that was repeated many times (**Learned**). We used the average time between entering two consecutive targets (inter-target interval, ITI) as a measure of performance.

**Figure 1.**
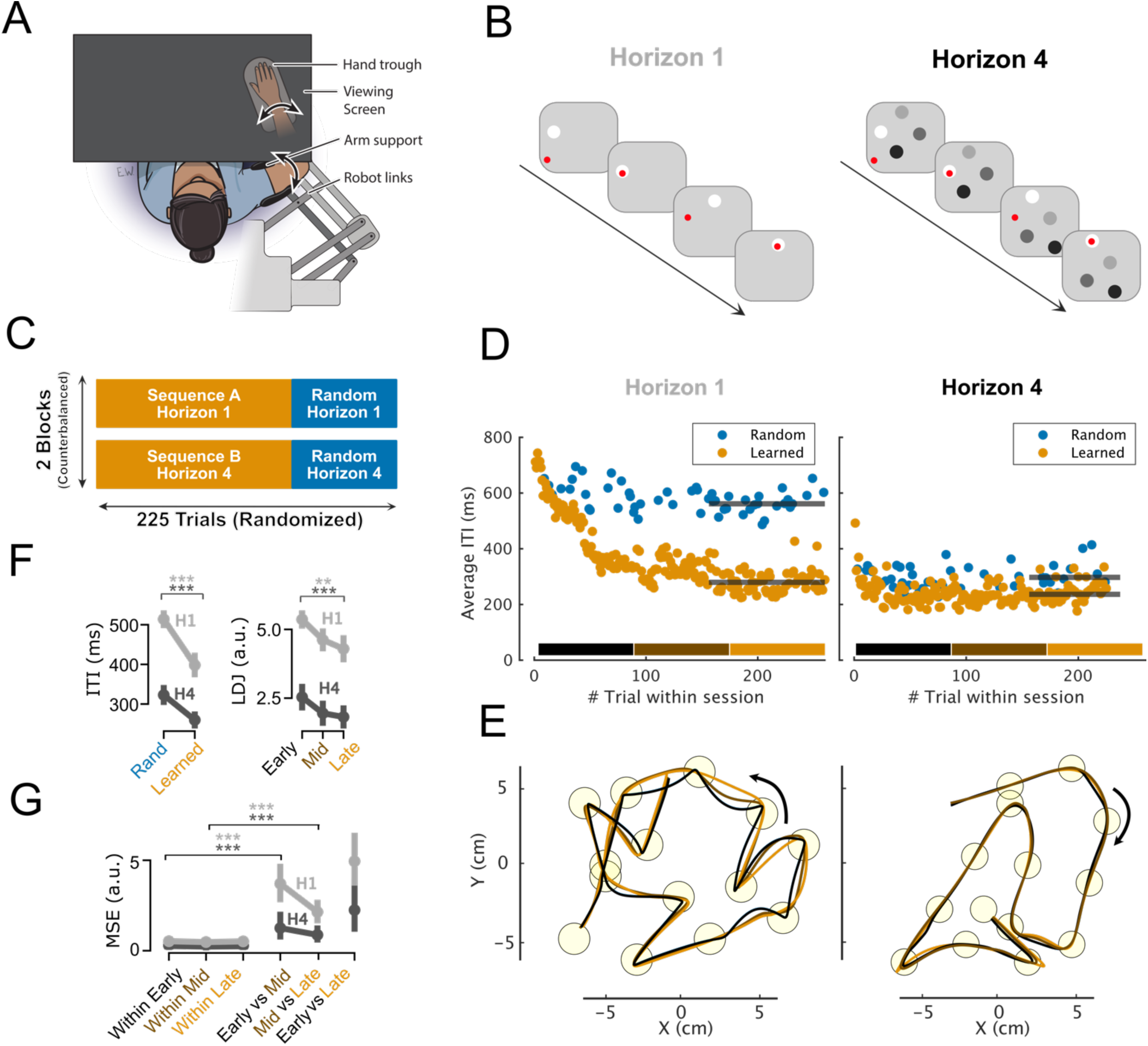
Learning sequences in Horizon 1 vs Horizon 4. **A)** Participants performed the task in an Exoskeleton Robot. A red cursor veridically indicated the tip of the index finger location. **B)** A cartoon example of targets shown to participants in Horizon 1 (left) and Horizon 4 (Right). The arrow shows time. For both Horizon 1 and Horizon 4, the task included: reach to the brightest target (box 1 to 2), dwell in the target for 50 ms, capture the target, the next target appears (box 2 to 3), reach to the new target (box 3 to 4). In Horizon 4, (left), in addition to the brightest target, 3 future targets are also shown on the screen, the order of these targets is shown with changes in brightness. **C)** Experimental design. Participants went through two blocks. In both blocks, they practiced one specific sequence by repeating it 180 times combined with 40 random sequences. In one block these sequences were in Horizon 1 and the other Horizon 4. The order of blocks was randomized for participants. **D)** Average Inter-target interval (ITI) in Horizon 1 (left), and Horizon 4 (right) for one representative participant. Orange and blue dots show average ITI for a learned or a random sequence trial respectively. Trials in the block are divided into three learning segments: Early (black), Mid (brown), and Late (orange). The horizontal black lines show the median ITI at the Late stage. **E)** Reach trajectories in Horizon 1 (left), and Horizon 4 (right) for one representative participant. Yellow circles show the targets of the sequence. Each trace shows participants average trajectories in Early (black), Mid (brown), and Late (orange) stages of the learning. The black arrow shows the direction movement. **F)** Left, Participants’ (n=14) speed, as measured by average Inter target interval (ITI). Right, participants’ movement smoothness as measured by average log dimensionless jerk (LDJ) in Horizon 1 (gray), and Horizon 4 (black) **G)** Participants’ trajectory changes during the learning process. Average dissimilarity (MSE) of trajectories is estimated between either withing or between three stages of learning (early, mid, late). For all plots: error bars show standard error. ** and *** show p < 0.01, p < 0.001.

### Learning in SRTT is mainly due to anticipation of the next target

In our first experiment, we assessed the role of anticipating future reach targets in sequence learning. We did so by asking participants (N=14, 8 Female) to practice a sequence either while they saw only one (Horizon 1) or four (Horizon 4) future reach targets. The Horizon 1 condition was similar to a standard serial reaction time task (SRTT) because the next reach target only appeared after the previous target was captured (**Fig. 1B, Horizon 1, Supplementary Video 1**). In the Horizon 1 condition, participants could become faster for learned sequences by anticipating the next target’s location and preparing their movement before it appeared. In the Horizon 4 condition, participants always saw four future reach targets so they could anticipate the next target’s location equally well for the random and learned sequences from the first trial **(Fig. 1B Horizon 4, Supplementary Video 2).** Hence, any difference between random and learned sequences in Horizon 4 should be unrelated to anticipating future reach targets.

Results for one representative participant are shown in **Figure 1D**. In Horizon 1, like many previous SRTT-like paradigms, the participant became faster (i.e. ITIs decreased) in executing the learned compared to the random sequence (**Fig. 1D, Horizon 1**). During the learning process, movement trajectories evolved: early in learning (black traces), trajectories displayed sharp transitions at each target, while later (orange traces), transitions became smoother (**Fig. 1E, Horizon 1**). Interestingly, when the same participant practiced a sequence with Horizon 4, they were immediately faster than in late stage of Horizon 1 (**Fig. 1E, Horizon 4**). Also, movement trajectories for Horizon 4 were smooth from the first trial, with only small adjustments over the course of learning (**Fig. 1E, Horizon 4**).

These observations were corroborated by the group-level statistics: practicing a sequence in Horizon 1 reduced ITIs by 115 ms compared to random sequences (t_(13)_ = 5.48, p=1.05e-4, d=1.60; Fig. 1F). Executing sequences in Horizon 4 was overall faster, even when comparing random sequences in Horizon 4 to fully practiced sequences in Horizon 1 (t_(13)_ = 3.34, p=5.34e-3, d=0.98). Faster movement production was also associated with smoother movements. We quantified movement smoothness by comparing the average Log Dimensionless Jerk (LDJ) in the early, middle, and late stages of learning (**Fig. 1F right, see Methods)**. Movements became smoother when comparing early to late learning stages in Horizon 1 (t_(13)_ = 3.59, p=3.26e-3, d=0.91). With the full ability to anticipate in Horizon 4, movements were overall smoother compared to Horizon 1 (F_(1, 13)_ = 107.33, p < 1.18e-7), even when comparing late Horizon 1 to early Horizon 4 (t_(13)_ = 4.86, p=3.12e-4, d=0.56). Overall, these results suggest that in the Horizon 1 condition participants mainly learned the to anticipate the next target’s location, allowing them to initiate movements faster and to curve the movements in ways that optimize the sequential trajectory. When the future targets were presented in the Horizon 4 conditions, participants showed the same behavior even for random sequences.

Critically, however, participants improved their ITI and movement smoothness with practice for the learned sequence even in the Horizon 4 condition. After practice they moved faster (Random vs Learned: t_(13)_ = 9.07, p=5.48e-7, d=1.06) and smoother (Early vs Late: t_(13)_ = 6.42, p=2.20e-5, d=0.55). This suggests that there were some improvements in sequence performance that did not depend on better anticipation of the next targets. One explanation for such improvements is that participants were able to specifically optimize the trajectory for the trained sequence. The continuous nature of the task allowed us to compare movement trajectories during the learning process (**Fig. 1E, Horizon 1 and 4**). To quantify these changes, we calculated the shape dissimilarity of movement trajectories in early trials to those of the mid and late trials (**see Methods**). We used the average dissimilarity of the trial within each stage as a baseline for comparison (**Fig. 1G**). In both Horizons, the trajectory dissimilarities between early versus mid (H1: t_(13)_ = 6.47, p=2.10e-5, d=2.31, H4: t_(13)_ = 3.22, p=6.61e-3, d=1.16) and mid versus late (H1: t_(13)_ = 7.99, p=2.00e-5, d=2.45, H4: t_(13)_ = 3.85, p=6.61e-3, d=1.43) were larger than their within learning stage baselines. This clearly shows that there was a gradual change in trajectory shape with learning, even in the Horizon 4 condition.

Together, these results show that in SRTT-like paradigms, most of the sequence learning reflects better anticipation of the future reach target, i.e. knowing *what* to do. Participants were immediately better if that knowledge was visually available to them (Horizon 4, see also Wong et al., 2015). Knowing what comes next also allowed participants to optimize movements for the sequence from the first trial (Kashefi et al., 2024). Importantly, however, even with the full knowledge of the future items, practice still led to improvement, suggesting that part of sequence learning can also reflect the optimization of specific movement trajectories, i.e. learning *how* to perform a sequence.

### Sequence learning with Horizon is effector specific

Learning in SRTT (Horizon 1) appears to be mostly related to anticipating the next target. Consistent with this idea, learning in the SRTT tends to transfer well from one hand to the other (Grafton et al., 2002; Verwey and Wright, 2004). Our results now suggest that participants also adjust their movement trajectories to optimize performance for the practiced sequence. Given that the refinement of the movement trajectories depends on the limb geometry and dynamics (Sainburg et al., 2003; Sainburg and Kalakanis, 2000), we would expect that there is little to no transfer between arms for this type of learning. Thus, by testing whether learning a sequence with one arm transfers to the other arm, we can obtain a measure about the degree to which the improvements are due to better anticipation of the target in allocentric space versus optimization of limb-specific movement trajectories.

In our second experiment, participants (N=14, 7F) completed a learning block and a probe block (**Fig. 2A**) both with Horizon 4. During the learning block, participants learned one sequence with their right hand and another with their left hand. These hand-specific sequences were practiced sixty times along with random sequences per hand to assess general learning. Trials were randomized across hands and sequence types (**see Methods**). By mixing left- and right-hand training, we ensured that both hands received general sequence training before the probe block.

**Figure 2.**
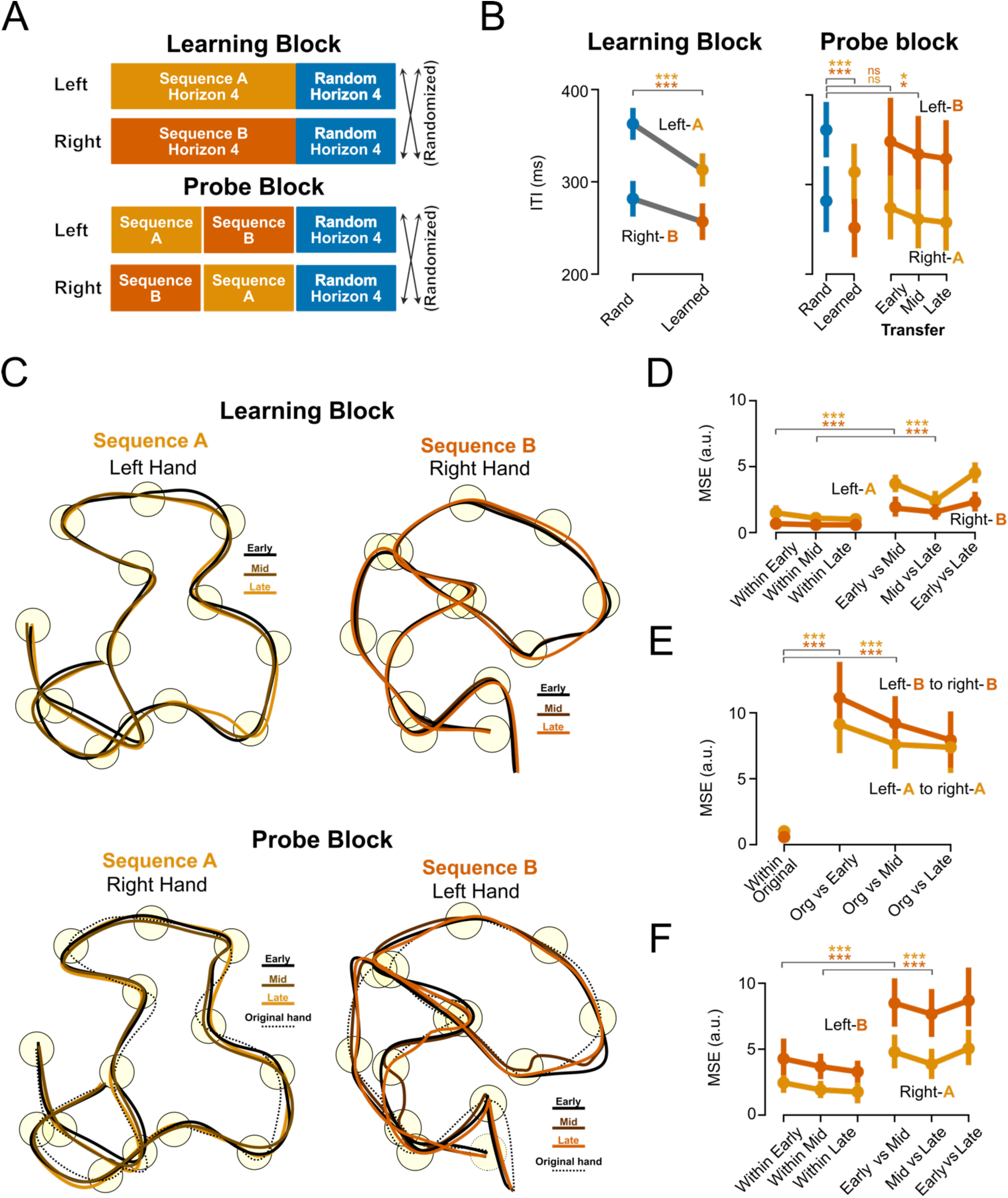
Learning with Horizon does not transfer between effectors. **A)** Experimental design. In the learning block, participants practiced one sequence with their right and another with their left hand, as well as a set of random sequences for each hand. Sequence type and hand were randomized. In the probe block, we tested the participants on three sequence types for each hand: A set of new random sequences, the sequence practiced with the same hand, and the sequence practiced with the opposite hand. **B)** Inter-target interval (ITI) for learning and probe blocks. Sequence A (orange) and B (red) are learned with the left and right hand respectively. In the probe block, Sequence A is also executed with the right hand and sequence B with the left. **C)** Average movement trajectories for one representative participant. Top: average trajectory in early, mid, and late stages of learning sequence A in the left hand and for sequence B with right hand. Bottom: average trajectories for sequence A and B when executed with the opposite hand (right and left) for the first 10 (early), second 10 (mid) and third 10 (late) trials. The black dotted trace shows how each sequence was executed in the late stages with the other hand. **D)** Changes in participants’ movement trajectory. Average dissimilarity of trajectories within or between three stages of learning (Early, mid, late) for sequence A (orange) or B (red). **E)** Trajectory dissimilarity between late stages of learning to early, mid and late stages of probe block, where the same sequence is executed with the opposite hand. **F)** Trajectory dissimilarity within or between early, mid, and late stages of executing the same sequences with the opposite hand in the probe block. For all plots: error bars show standard error. ** and *** show p < 0.01, p < 0.001.

By the end of the learning block, participants’ ITI for the practiced sequence had significantly decreased for both the left (t_(13)_ = 7.52, p=4.00e-6, d=0.91) and right (t_(13)_ = 5.372, p=1.27e-4, d=0.40) hands. As expected, the left hand was generally slower since all participants were right-handed (F_(1, 13)_ = 82.61, p < 5.38e-7; **Fig. 2B, Learning Block**). In the probe block, participants were tested on a new set of random sequences (20 trials per hand), the sequence learned with the same hand (20 trials), and the sequence learned with the opposite hand (30 trials). The learned sequences were transferred in extrinsic coordinates, meaning that all visual cues remained consistent between learning and probe blocks. As in the end of learning, participants continued to perform slower on the random sequences than on the sequences learned with that hand (**Fig. 2B, Probe Block left**). Importantly, the first 10 trials of the transferred sequence were not significantly different from random sequence (left hand: t_(13)_ = 1.10, p=0.280; right hand: t_(13)_ = 1.41, p=0.18). It was only after another 10 trials that the ITIs became significantly different from the random sequences (Left hand: t_(13)_ = 3.0, p=1.03e-2, d=0.4; Right hand: t_(13)_ = 4.17, p=1.09e-3, d=0.32), indicating that participants started to learn the transferred sequence with the new hand.

We also monitored changes in movement trajectories in both blocks. **Figure 2C** shows average trajectories during learning for one representative participant. During the initial learning, similar to the first experiment, movement trajectories for both hands changed during learning (**Fig. 2C, Learning block**). To quantify this observation at the group level, we again used trajectory dissimilarity between the early, mid, and late stages of learning and compared these to the dissimilarity within each of these stages as a baseline. In the learning block, we replicated Experiment 1 with two hands. The dissimilarity between early versus mid (Left-A: t_(13)_ = 13.28, p=6.14e-9, d=2.71, Right-B: t_(13)_ = 5.142, p=1.16e-4, d=1.52) and mid versus late (Left-A: t_(13)_ = 7.14, p=8.00e-6, d=1.69, Right-B: t_(13)_ = 6.70, p=1.50e-5, d=1.67) were both larger than their respective baselines, suggesting a systematic change in movement trajectories with learning (**Fig. 2D**).

Importantly, in line with our prediction, after the transfer to the other hand, participants executed the same sequence with a different trajectory (**Fig. 2C, Probe block)**. The trajectory for the early, mid and late stages of the probe block (solid traces) were qualitatively different from the trajectory from the late learning block (dotted traces). We quantified this by comparing the trajectories during late learning to that of the first (early), second (mid), and third (late) 10 trials with the trajectory from the opposite hand (**Fig. 2E**). For both hands, the early transfer trials were highly dissimilar to how the sequence was executed in the late learning block. The same was true for middle and late transfer trials, which were also highly dissimilar to how they were executed originally in the learning block (In both hands, all learning stage comparisons: t_(13)_>6.52, p< 1.90e-5). These results show that participants had to acquire a new trajectory to execute the same sequence with the other hand. After the transfer, the trajectories kept changing within the new effector. The dissimilarity between post-transfer early versus mid trials was higher than within early baseline (Right-A: t_(13)_ = 6.11, p=3.70e-5, d=1.37, Left-B: t_(13)_ = 7.41, p=5.00e-6, d=1.52). The same was true for mid versus late dissimilarities (Right-A: t_(13)_ = 5.50, p=1.00e-4, d=1.32, Left-B: t_(13)_ = 6.79, p=1.30e-5, d=1.66) (**Fig. 2F**). Together, the post-transfer ITI results and trajectory changes show that participants had to relearn a trajectory optimized for the new effector.

The ITI results from this experiment also confirm that learning with the Horizon of future targets was not due to enhanced anticipation of future targets. Since the visual cues in both the original and transferred sequences were identical, any improvements in anticipation should have been immediately transferable to the other effector—an effect we did not observe. Instead, the trajectory data suggest that learning with Horizon is associated with effector-specific modifications in movement trajectories, occurring only for the effector with which the sequence was originally learned.

### Sequence learning with horizon is not due to improvements in single reaches or their transitions

Our experiments so far have shown that practicing a sequence in Horizon 4 leads to fine-tuning of hand- and sequence-specific trajectories. How are these trajectories represented? At one extreme, the motor system may store the entire trajectory as one single unit. At the other extreme, the motor system may have optimized small chunks of the sequence, for example a curved movement through three targets. To answer this question, we tested how many elements of the original sequence are necessary to trigger the learned ITI reduction. Answering this question could provide insight into the fundamental unit of sequence learning - individual reaches, transitioning between reaches, or potentially a chunk consisting of multiple reaches and their transitions.

We conducted a two-block experiment (**Fig. 3A**). In both blocks sequences were presented in Horizon 4. In the first block, participants (N=28, 16F) practiced a single subject-specific sequence for 160 trials, along with 40 random sequences. In the second block, we randomly selected between 1 and 5 consecutive reach targets from the practiced sequence (orange circles, dotted line) and embedded them at random locations in otherwise random sequences (blue circles solid lines) (**Fig. 3B, See Methods**). Embedding a single target tests for improvements in reaching a specific target regardless of start location (**Fig. 3B, 1 Target**). Embedding two targets tests for improvements in executing individual reaches. Embedding three targets tests for improvements in transitions between reaches (**Fig. 3B, 3 Targets**), while embedding more than three targets assesses performance on larger chunks of the sequence, involving multiple reaches and transitions (**Fig. 3B, 5 Targets**).

**Figure 3.**
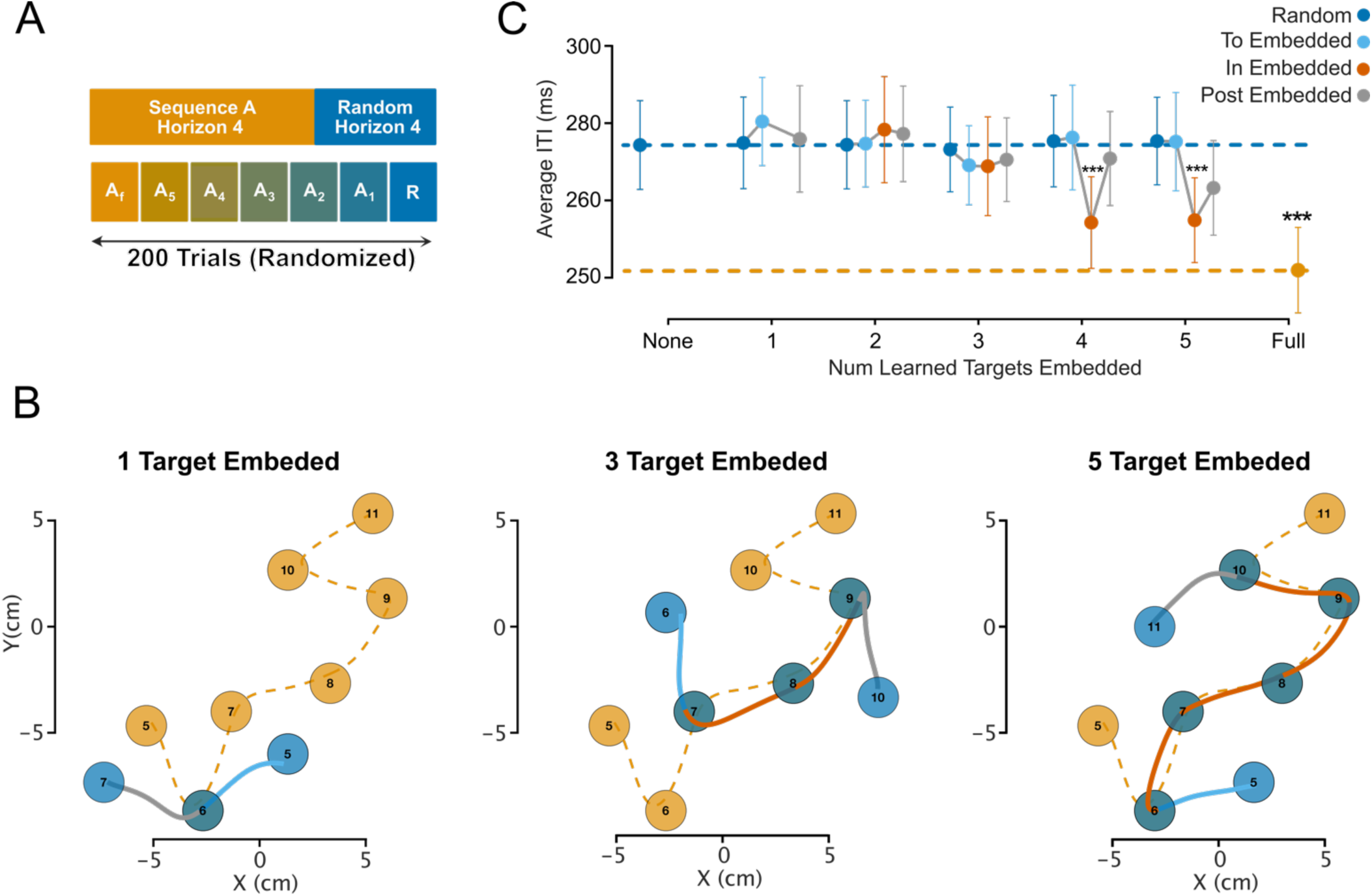
Learning with horizon is not due to improvements in single reaches or transition of reaches. **A)** Experimental Design. Participants completed one block with 160 trials of a specific sequence, interspersed with 40 random sequences. In the second block, participants performed fully random sequences (R), fully practiced sequences (A_f_), or sequences containing 5 to 1 practiced reach targets embedded within otherwise random sequences (A_5_ to A_1_). **B)** Example embedded trials. Orange targets: targets 5 to 11 of the practiced reach sequence for one representative participant. Blue targets: random sequences with one, three, or five embedded targets from the practiced sequence. **C)** Average ITI for Block 2 for random reaches (blue), reaches before (light blue), in (red) and after (gray) the embedded sequence. Error bars show the standard error of the mean, and *** indicates p < 0.001.

The participants’ average ITI during fully random (dotted blue line) and fully practiced (dotted orange line) trials in Block 2 served as upper and lower reference, respectively (**Fig. 3C, see Methods**). In the trials with embedded parts of the trained sequences, the ITIs did not differ significantly from fully random sequences, neither before or after the embedded sections (**Fig. 3C, dark blue dots**). Additionally, we observed no speed-up in reaches leading into (t_(27)_ < 1.45, p>0.15 for all num targets embedded) (**Fig. 3C, light blue dots**) or exiting (t_(27)_ < 2.01, p>0.054 for all num targets embedded) (**Fig. 3C, gray dots**) the embedded segments. A reduction in ITI was only observed for reaches when the embedded section contained four (t_(27)_ = 6.16, p=1.31e-6, d=0.33) or five targets (t_(27)_ = 5.27, p=1.48e-5, d=0.34). No speed-up occurred when only two or three (t_(27)_ < 1) targets were embedded.

This experiment revealed that the learned improvements did not occur at the level of single targets, single reaches, or even transitions between reaches. Instead, the improvements emerged over a longer time scale and are only triggered when a larger chunk of the practiced sequence is required.

### Learning with Horizon is prompted by the context of previous reaches

We then asked where within the embedded segment the speed-up occurred. The pattern of ITI reduction can reveal whether the history of the previous reaches or the look ahead of future reaches is required to trigger the sequence-specific memory. If the previous reaches are essential, we would expect less ITI reduction at the beginning of the embedded segment, since the first embedded reach is preceded by random reaches. Similarly, if anticipation of future reaches is important, we would expect smaller ITI reduction toward the later part of the embedded segment since the last embedded reach is followed by random reaches.

Within the embedded section (**Fig. 4A, B**), the ITI for the first reach (green dot) was not significantly different from that of the reaches in random sequences. This was true regardless of how many targets were embedded (t_(27)_ < 2 for all num targets embedded). Reliable ITI reductions were observed only for the second, third, and fourth reaches in the embedded section for trials with four targets embedded (second: t_(27)_ = 3.45, p=1.81e-3, d=0.38; third: t_(27)_ = 3.57, p=1.34e-3, d=0.35). The same was true for trials with five targets embedded (second: t_(27)_ = 3.09, p=4.60e-3, d=0.41; third: t_(27)_ = 2.37, p=2.49e-2, d=0.24; fourth: t_(27)_ = 3.36, p=2.32e-3, d=0.39). The lack of speed-up in the first embedded reach clearly shows the importance of the immediate history of reaches.

**Figure 4.**
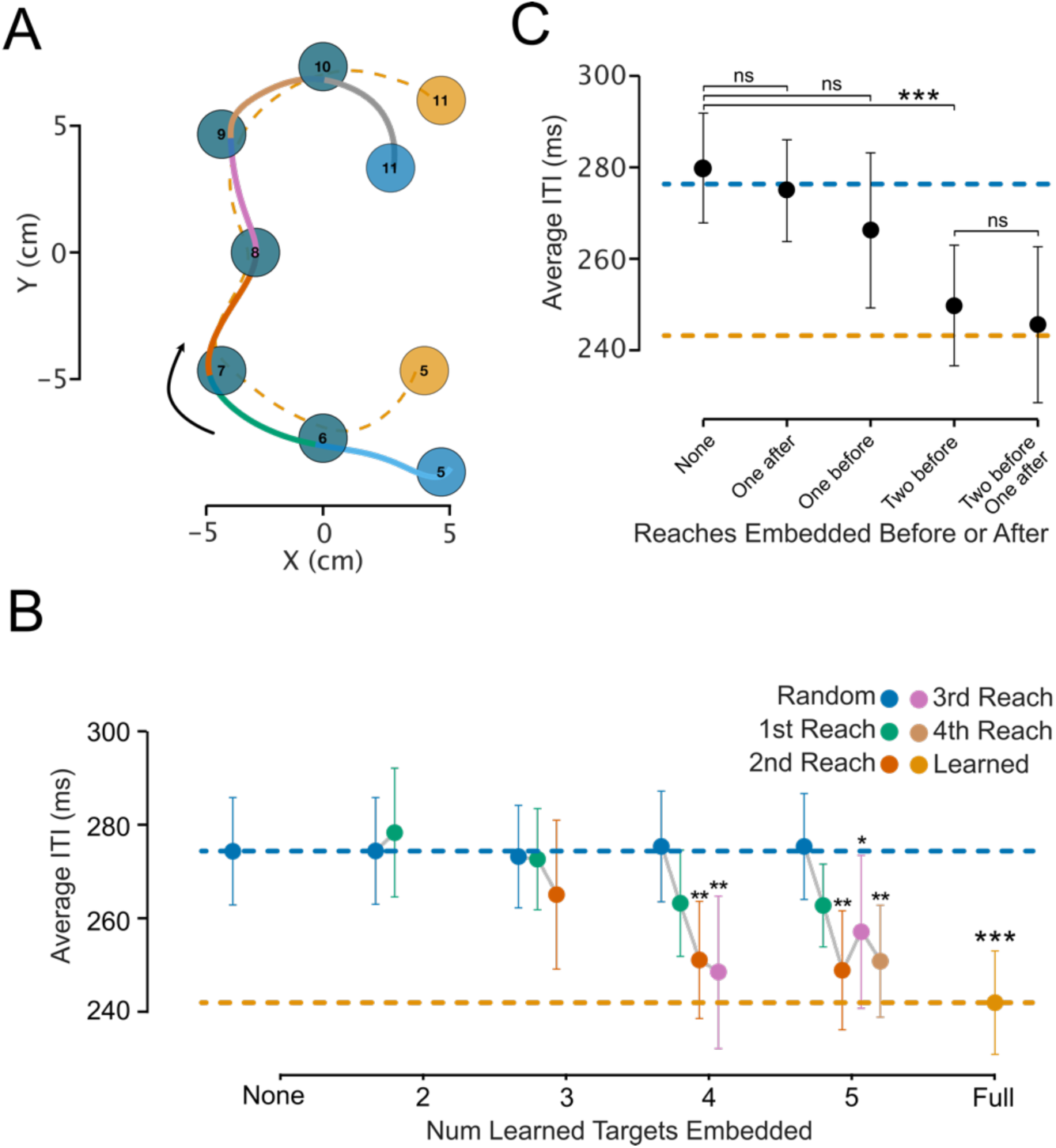
Learning with horizon is context dependent. **A)** Sample trial from a 5-target embedded sequence. Orange circles: targets 5 to 11 of the practiced sequence. Blue circles: targets 5 to 11 for the embedded trial, with reaches leading into (light blue) and exiting (gray) the embedded section. **B)** Average inter-target interval (ITI) for embedded trials. The blue and orange dotted lines represent ITIs for fully random and fully practiced trials, respectively. Dark blue dots correspond to ITIs for random reaches outside the embedded segments in the embedded trials. As in A, the green, red, pink, and beige dots indicate ITIs for the four possible reaches within the embedded section. **C)** Average inter-target interval (ITI) for practiced reaches without practice reaches before or after (None), with one reach following (One After), one reach preceding (One Before), two reaches preceding (Two Before), or two preceding and one following (Two Before, One after). Error bars show the standard error of the mean, and *, **, and *** indicates p < 0.05, p < 0.01, and p < 0.001.

To summarize the importance of past and future reaches across trials with different embedding length, we used isolated embedded reaches as a baseline (**Fig. 4C, None**). Relative to this, we found no ITI speed-up when only one future came from the practiced sequence (t_(27)_ = 0.7, p = 0.48), suggesting that anticipating a single trained reach is insufficient to trigger speed-up (**Fig. 4, One After**). When a single trained reach preceded the current reach, we observed a non-significant trend toward reduced ITI (t_(27)_ = 2.04, p = 0.0504, d = 0.173). A significant ITI reduction only occurred when two preceding reaches were trained (t_(27)_ = 4.93, p = 3.60e-5, d = 0.44), highlighting the importance of preceding reach context in triggering the speed-up. Finally, embedded reaches flanked by two embedded reaches before and one after showed a small, non-significant additional reduction compared to two preceding embedded reaches alone (t_(27)_ = 0.568, p = 0.57), reinforcing that future reach context has minimal impact on ITI reduction. These findings underscore that what is learned in the Horizon task is only activated in the context from two familiar previous reaches.

## Discussion

The mechanisms that lead to motor sequence learning can be separated into three broad categories. First, improvements can occur because participants become better at selecting and executing each individual movement, independent of their specific sequential context. Second, participants can learn to anticipate *what* movement to make next, thereby initiating the movement more quickly and/or preparing it better. Third, participants can develop a sequence-specific representation that encodes *how* each movement should be performed within the context of the sequence. In this paper we sought to determine to what degree each of the mechanisms can explain improvements in a continuous reaching task. First, we compared performance in trained sequences with that in random sequences to ascertain that the improvements were not due to improvements in executing isolated movements. By showing the participants either only the next target (Horizon 1), or the next four targets (Horizon 4), we then contrasted learning with and without information about *what* comes next.

Being able to see the future reach targets accelerated the participant’s performance: even in the first trial in Horizon 4, they moved faster and more smoothly than in the late stages of learning in Horizon 1. This observation fully supports the conclusions drawn by Wong et. al. (2015), who demonstrated that in a discrete serial reaction time task (SRTT), most of the decrease in movement time is due to participants learning to anticipate the future targets. Our results also show that knowing “what” to do not only speeds up the initiation of the movement – but also allowed participants to co-articulate the movements in anticipation of the next targets, leading to much smoother trajectories.

In contrast to Wong et. al. (2015), however, we also found clear evidence for sequence-specific improvements with practice even when participants had full knowledge (i.e. could see) of the future targets (in Horizon 4). Participants achieved this improvement by optimizing *how* movements were executed in the sequence, evidenced by a gradual change of the trajectory across the learning process. We then showed that this learned sequence representation is effector specific and that it does represent a sequential context of three or more reaches.

### Learning with Horizon cannot be due to motivational factors

The sequence-specific improvements observed in Horizon 4 might be attributed to enhanced movement vigor. Confidence and familiarity gained through practicing a sequence in Horizon 4 could function as an internal reward, driving a general motivational boost and leading to more vigorous movements. Wong et al. (2015) previously argued for such a mechanism as one factor leading to better performance for trained sequences. In their experiment, participants moved faster for an entire sequence, even if only the beginning of the sequence was known. In our case, such motivational factors are unlikely to explain the observed advantage over random sequences. First, if familiarity of the practiced sequence increased the motivation of the participants, we should also have found faster performance when the same sequence was performed with the other arm, which we did not observe. Second, speed-up due to motivational factors should have transferred to unfamiliar segments following familiar segments, which we also did not observe.

### Learning what to do next

Our results support the conclusion that improvements in the SRTT task are mainly due to learning what the next movement will be (Howard et al., 1992; Wong et al., 2015). SRRT tasks became a popular paradigm to study sequence learning, because it was shown that learning in these tasks can occur outside conscious awareness, and independent of episodic memory systems. This suggested that the SRTT may rely on a single procedural memory system, unrelated to conscience anticipation of what should be done next (Nissen and Bullemer, 1987). However, there are at least three pieces of evidence suggesting that sequence-specific improvements observed in SRTT can nevertheless be influenced by anticipatory processes, whether these are accessible to consciousness or not. First, after learning, participants demonstrate some fragmentary knowledge of sequence cues when probed with sensitive measures like recognition or production tasks (Reber and Squire, 1998); this observation is true even in amnestic patients (Reber and Squire, 1994). Most reported implicit learning in SRTT is due to participants learning fragmentary knowledge of the sequence items by learning the distribution of presented cues (for review, see Shanks, 2005; Shanks and St. John, 1994). Second, learned improvements often transfer to other effectors either fully or partially (Cohen et al., 1990; Grafton et al., 2002; Verwey and Wright, 2004). Third, when a task is designed such that participants can fully predict the next cues, practicing the sequence does not lead to any sequence-specific improvements (Howard et al., 1992; Wong et al., 2015).

To study motoric aspects of sequence learning, we provided participants with clear and direct knowledge of the upcoming sequence items in our Horizon 4 condition. This is common practice in the discrete sequence production (DSP) task, in which participants are presented with an entire sequence and are instructed to execute it as fast as possible. Even under such conditions, however, learning improvements cannot be attributed unambiguously to pure *motor learning*. For example, in most finger motor sequence task, the required finger presses are indicated as a list of digits (Berlot et al., 2020; Karni et al., 1995; Rhodes et al., 2004). While the digits provide complete information about the sequence items, the stimulus-response mapping from numbers to fingers is relatively abstract and requires a time-consuming process. Consequently, a substantial portion of learning in these tasks can be explained by an improved ability to perform the process of stimulus-response mapping quickly and in parallel with ongoing execution (Ariani et al., 2024, 2021; Kornysheva and Diedrichsen, 2014; Shahbazi et al., 2024a). Consistent with this idea, learning of DSP tasks often generalizes at least partially to other effectors (Wiestler et al., 2014; see Abrahamse et al. 2013 for a review).

In contrast, in our reaching task, learning was nearly entirely limb specific. One possible explanation for this difference is that the stimulus response-mapping from spatially presented targets and reaching movements towards these targets is already very fast and automatic (Day and Lyon, 2000; Diedrichsen et al., 2004, 2001; Goodale and Milner, 1992; Pruszynski et al., 2010), such that sequence learning cannot further improve the speed of the translation from the target to the desired movement.

### Learning how to perform a sequence: representation and possible neuronal mechanisms

In Horizon 4, we observed effector-dependent, sequence-specific improvements. Our embedding experiment (**Fig. 3, 4**) demonstrated that more than 3 trained reaches are needed to see a behavioral advantage. This shows that learning did not occur at the level of single or even pairs of reaches. Instead, the learned representations encapsulate a longer sequential context of the movement, without being a fixed representation of the entire sequence (Buchner et al., 1998; Shahbazi et al., 2024b).

These results suggest that participants planned a continuous trajectory through the multiple targets ahead (Wong et al., 2016). Over the course of learning, repeated execution of this trajectory led to gradual improvements in the trajectory’s precision and efficiency (Shmuelof et al., 2012). When encountering the same targets within a random sequence (**Fig. 3 and Fig. 4**), participants could recall a motor memory of how to execute thus previously practiced trajectory shapes to accelerate movement execution (Morasso and Mussa Ivaldi, 1982; Viviani and Terzuolo, 1982). An unresolved question, however, remains whether participants learned this sub-sequence representation in our task by chunking the sequence items into discrete segments (Ramkumar et al., 2016; Verwey, 1996), or whether trajectory planning is a continuous process that is updated with every new target. In our embedding experiment we picked segments from the learned sequence at random, such that these segments would have only sometimes aligned with learned “chunks”. Nevertheless, we observed a relatively consistent acceleration of the movement whenever a segment was repeated, making a target-by-target *continuous* planning hypothesis seem more likely. Nonetheless, we acknowledge our experiment does not provide a strong test of the *chunked* versus *continuous* sequence planning hypotheses - indeed these two ideas are difficult to distinguish using behavioral experiments only. We believe, however, that the two ideas make different predictions about the neural implementation.

The relatively long duration of the learned elements makes it unlikely that the underlying representations are learned in primary motor cortex (M1). Previous studies have shown that activity patterns in M1 can be explained by a superposition of the activity related to the individual movements, both for reaches (Zimnik and Churchland, 2021) and finger presses (Yokoi et al., 2018) – with very little evidence that the representation of individual movement elements changes with the sequential context. Thus, it is possible that M1 is blind to sequential dependencies at least over the time window identified in our experiments. Learning long sequence of movement has been reported to be associated with changes in parietal and premotor areas (Berlot et al., 2021, 2020). These areas likely take an intermediate position between deciding *what* to do next and *how* to do it (Diedrichsen and Kornysheva, 2015; Wong et al., 2016). That is, these areas may not fully encode all the motoric details of the movement but may seed an effector-specific region in M1, which allows it to execute the movements with an optimized trajectory (Churchland et al., 2010).

At the level of neural dynamics in execution related areas, sequence-specific optimization of individual movements could potentially be implemented in neuronal populations by selecting the optimal preparatory state for each reach (**Fig. 5**). In Horizon 1, since no information about the upcoming movements is known, the preparatory state for reaching to target A is selected among a large set of potential initial states that all lead to movement to target A, leading to natural trial to trial variability of reaches. In the case of Horizon 4 and before any practice (Random), information about the next target cue is used to seed the system with a preparatory state that leads to biases in reach kinematics that are optimized for the next reach in the sequence. With practice of a sequence in Horizon 4, the sequence representation formed in other brain areas allows further refinement of the preparatory states that leads to reaches optimized for even longer segments of the sequence. Alternatively, producing a learned sequence might involve preparing chunks of multiple movements as a single long movement in higher motor areas, which are then sent to effector-specific regions like M1 for execution. At the neural level, *chunked* and *continuous* hypotheses of sequence planning can be distinguished by examining neural states at chunk boundaries. Under the *chunked* hypothesis, neural states should undergo larger transitions between chunks compared to within-chunk movements. Conversely, the *continuous* planning hypothesis predicts no major transitions at chunk boundaries, as movements are always prepared individually. Future electrophysiological experiments are necessary to test these potential mechanisms. The continuous nature of our sequence paradigm offers a useful framework for investigating these hypotheses.

**Figure 5.**
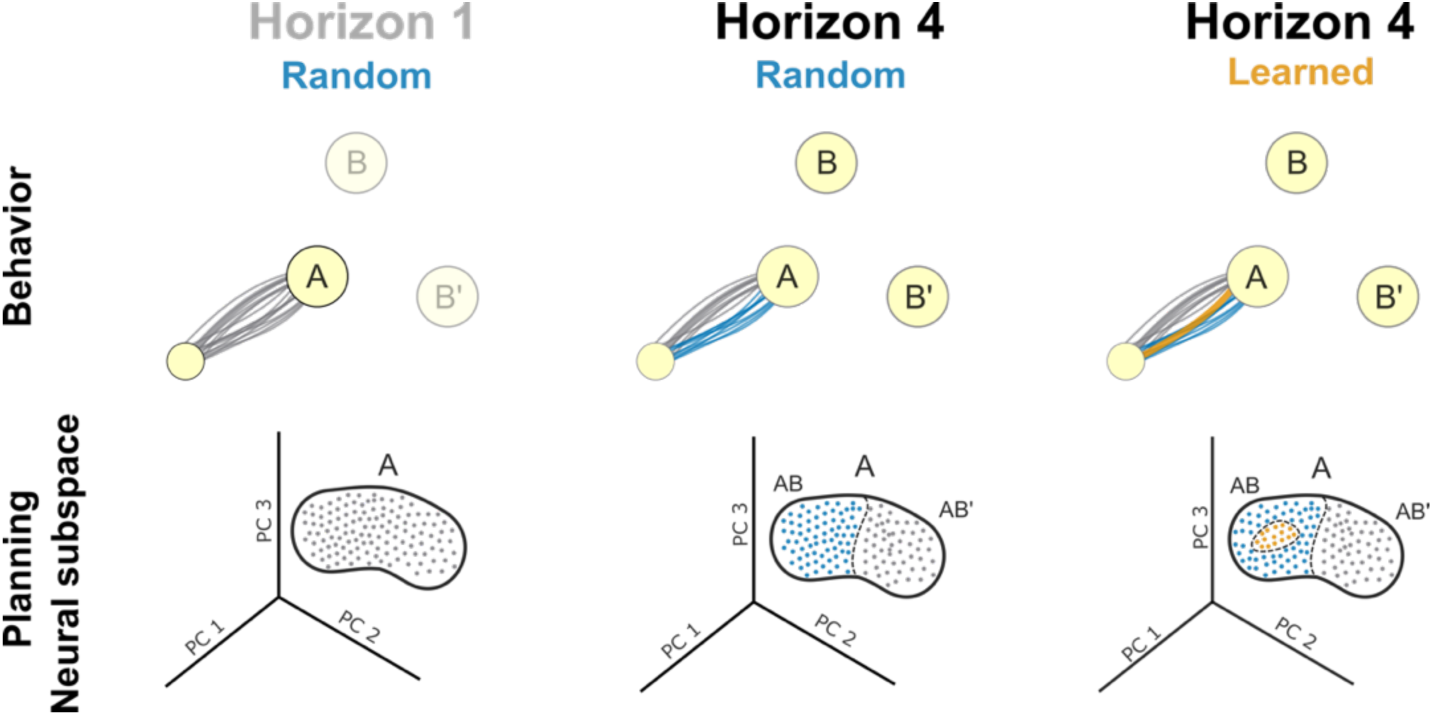
A potential neural implementation of sequence learning with Horizon. The first row shows reaching to target A in Horizon 1, with no target (B or B’) shown on the screen, Horizon 4 before any practice, and Horizon 4 when sequence is practiced. Second row shows the planning subspace for all the conditions shown in first row.

## Methods

### Participants

Our study consisted of three experiments. For Experiment 1, we recruited 14 participants (8 female) with an average age of 22.4 ± 5.4 years (**Fig. 1**). All were right-handed, with an average handedness score of 68 ± 24, assessed using the Edinburgh Handedness Inventory (Oldfield, 1971). For Experiment 2 (**Fig. 2**), we recruited an additional 14 participants (7 female) with an average age of 24.3 ± 3.4 years and an average handedness score of 72 ± 22. For Experiment 3 (**Fig. 3** and **4**), we recruited 28 participants (14 female), with an average age of 22.9 ± 4.1 years and an average handedness score of 69 ± 30. All participants reported no history of musculoskeletal, neurological, or psychiatric disorders. Data collection for each experiment took place within a single session. At the start of each session, participants provided informed consent and completed the Edinburgh Handedness Inventory. Participants were compensated for their time (CA$15 per hour). All procedures were approved by the Health Sciences Research Ethics Board at the University of Western Ontario.

### Apparatus

Participants performed all trials in the KINARM exoskeleton robot (Scott, 1999). Participants rested their right arm (or both arms in bimanual experiments) in the robot while being seated in a height-adjustable chair. The robot supported the weight of the arm, allowing free movement of the hand in the horizontal plane. Reach targets were displayed in the plane of the task via a virtual display, which blocked the participants’ direct view of their arms. The veridical position of their fingertip was displayed on the display as a circular cursor (0.5 cm diameter); (**Fig. 1A**). Hand kinematics were recorded at a sampling rate of 1000 Hz.

### General procedures

In all three experiments, participants moved the circular cursor to 14 circular targets (1 cm diameter) to complete each trial. The reach targets were randomly placed within a 10 x 10 cm workspace, following two rules: (1) the distance between consecutive target centers was set between 4.8 and 5 cm, ensuring consistent reach distances; (2) the minimum distance between any four consecutive targets was at least 3 cm to prevent overlap when multiple targets were displayed. Using these criteria, a unique set of sequences was generated for each participant. At the beginning of each trial, the first target appeared as a home position, and participants moved the cursor into this target and waited for a go-cue. After 300 ms, depending on the Horizon condition, either 1 (Horizon 1) or 4 (Horizon 4) upcoming targets were displayed. Target order was indicated by brightness, with the first target as the brightest. Following an additional delay of 300–500 ms, the disappearance of the home target served as the go-cue, prompting participants to reach the brightest target as quickly and accurately as possible. For each target, participants were required to “capture” it by dwelling inside it for at least 50 ms. Once captured, the target disappeared, the remaining targets’ brightness updated, and a new target appeared with the lowest brightness. This process repeated until all 14 targets in the sequence were completed (**Fig. 1B, Supplementary Video 2**). Trials were interrupted with an error message under three conditions: (1) if the participant left the home target before the go cue, (2) if they exited a target before the 50 ms dwell time, or (3) if their reach time exceeded 500 ms for any of the 14 targets. Interrupted trials were repeated later in the block. The number of interrupted trials was < 5% in all participants across all the experiments.

### Procedures for Experiment 1

We evaluated participants’ learning with and without advanced knowledge of future targets (Horizon conditions). Each participant completed two blocks: one in Horizon 1 and the other in Horizon 4. For each block, we generated 46 unique sequences based on the rules described in the general procedures, then randomly designated one sequence as the “learning sequence.” This specific sequence was repeated 180 times to assess sequence-specific learning, while the remaining 45 sequences were each presented once to assess sequence-general learning. The trial order was randomized within each block. After a 5-minute break, participants performed the second block, which contained the same number of trials but under the alternate Horizon condition. The order of Horizon conditions was counterbalanced across participants (**Fig. 1C**).

### Procedures for Experiment 2

We tested whether sequence learning with Horizon 4 transfers across effectors. Participants completed a learning block and a probe block. For the learning block, we generated 31 unique sequences for each hand. One sequence was randomly designated to be repeated 60 times to assess sequence-specific learning, while the remaining 30 sequences were each presented once to assess general learning, resulting in a total of 90 trials per hand. Sequence type (learned or random) and hand (left or right) were randomized within the block (**Fig. 2A, Learning Block**). This design also ensured that both hands were trained in the general task procedure before the probe block. Each hand had its own cursor—a red cursor for the right hand and a blue cursor for the left hand—which accurately represented the participant’s index finger position. Only one cursor was activated per trial, instructing participants which hand to use. After a 5-minute break, participants moved on to the probe block to assess learning transfer. In this block, each hand completed three types of trials: 20 trials with new random sequences, 20 trials with the sequence learned by the same hand in the learning block, and critically, 30 trials with the sequence learned by the opposite hand in the learning block. Trial order, including hand and sequence type, was randomized within the probe block (**Fig. 2A, Probe Block**).

### Procedures for Experiment 3

We assessed generalization to random sequences that contained parts of trained sequence. We generated 41 unique sequences and randomly selected one to be repeated for 160 trials to evaluate sequence-specific learning, with the remaining 40 sequences presented only once (**Fig. 3A**). All sequences were presented in the Horizon 4 condition. After a 5-minute break, we tested generalization by introducing a set of modified sequences that shared 1 to 5 consecutive targets with the learned sequence (**Fig. 3B**). Apart from these embedded segments, the rest of each sequence was designed to resemble a random sequence. The embedded targets were randomly selected from the original sequence and placed in random locations within the new sequences, ensuring no bias from specific positions in the original sequence. Once the embedded targets were chosen, random targets were added before and after the embedded segment, following the rules explained in general procedures. Additionally, to prevent unintentional similarities with the originally learned sequence, we enforced an extra rule: the targets immediately before and after the embedded segment had to be at least 3 cm away from the corresponding targets in the original sequence. Following these rules, we created 30 trials for each embedding condition (1 to 5 embedded targets). The block also included 30 trials of the original learned sequence and 40 trials of entirely new random sequences (**Fig. 3A**).

### Time, smoothness, and trajectory change analysis

We used Inter-Target Interval (ITI) as a measure of participants’ movement time. ITI was defined as the time taken for participants to move the cursor from the boundary of one target to the next. For the first reach, since participants started from the center of the home target, ITI was calculated from the onset of the go cue to the moment they reached the boundary of the first target. We averaged 14 ITIs for 14 reaches in a trial to obtain a single movement time measure for each trial. In all experiments, we assessed learning by comparing the average ITIs for sequences that were repeated only once (random sequences) with those for one specific sequence repeated multiple times (learned sequence). This comparison allowed us to distinguish improvements specific to practicing a specific sequence (sequence-specific learning) from general improvements due to factors like increased familiarity with the apparatus or overall improvement in sequence execution (sequence-general learning).

We measured participants’ movement smoothness using Log Dimensionless Jerk (LDJ) (Balasubramanian et al., 2015). LDJ was chosen over simpler metrics, such as the sum of squared jerk, because it offers greater robustness when comparing movements with varying durations or peak speeds (Balasubramanian et al., 2012; Hogan and Sternad, 2009). We first divided each trial into 14 consecutive reaches, defined according to our ITI criteria, as movements from one target boundary to the next. We calculated LDJ separately for each reach and then averaged these values to obtain a single smoothness measure per trial.

To plot the average trajectories of the representative participant during the learning process (**Fig. 1E**, **Fig. 2C**), we first segmented each trial into 14 consecutive reaches. We then divided the trials within the learning block into three segments (early, mid, and late) to capture different learning stages. For each reach, we resampled the x and y hand positions to match the median trial length within each learning segment, ensuring an equal number of samples across trials. Finally, we averaged these resampled trajectories across trials, yielding representative hand trajectories for early, mid, and late stages of learning.

To quantify systematic changes in movement trajectories during learning, we divided the trials into three stages: early, mid, and late learning. We then assessed the dissimilarity between each pair of learning stages (e.g., early vs. mid), using within-segment dissimilarity as a baseline (e.g., early vs. early). To estimate dissimilarity between two stages, we randomly selected one trial from each stage and measured the difference using dynamic time warping (DTW) (Paliwal et al., 1982; Sakoe and Chiba, 1978). This random sampling and DTW calculation were repeated 5000 times, and we reported the average of these 5000 samples as the estimated dissimilarity between learning stages. To estimate within-stage dissimilarity, we applied the same method, sampling two trials from the same stage (without replacement) for each comparison.

### Statistical analysis

We used a within-subject design across all experiments. Statistical analyses were conducted using Statsmodels 0.14.4 (Seabold and Perktold, 2010). For each test, we report degrees of freedom, test statistics, and p-values in the text. We employed two-way repeated measures analysis of variance (ANOVA) and paired t-tests. In Experiment 1, the factors were Horizon (two levels: Horizon 1 and Horizon 4), sequence type (two levels: Random and Learned), or learning stage (three levels: early, mid, late). In Experiment 2, the factors included hand (two levels: left and right), sequence type (two levels: Random and Learned), or learning stage (three levels: early, mid, late). All t-tests were two-sided.

## Data and materials availability

All raw data generated in this study, along with the code for analysis and visualization, will be made publicly available upon publication.

## Authors contribution

<u>Mehrdad Kashefi</u>: Conceptualization, Data acquisition, Analysis, Visualization, Methodology, Writing – original draft, review and editing; <u>Jörn Diedrichsen</u>: Conceptualization, Supervision, Methodology, Writing – review and editing; <u>J. Andrew Pruszynski</u>: Conceptualization, Supervision, Resources, Methodology, Writing – review and editing.

## Acknowledgements

This work was supported by a CIHR Project Grant to JD and JAP (PJT-175010). JAP received a salary award from the Canada Research Chairs Program.

